# DECODING SECOND ORDER ISOMORPHISMS IN THE BRAIN: The case of colors and letters

**DOI:** 10.1101/2020.05.04.076703

**Authors:** Stephen José Hanson, Leyla Roskan Caglar, Catherine Hanson

## Abstract

We introduce a new method for decoding neural data from fMRI. It is based on two assumptions, first that neural representation is distributed over networks of neurons embeded in voxel noise and second that the stimuli can be decoded as learned relations from sets of categorical stimuli. We illustrate these principles with two types of stimuli, color (wavelength) and letters (visual shape), both of which have early visual system response, but at the same time must be learned within a given function or category (color contrast, alphabet). Key to the decoding method is reducing the stimulus cross-correlation by a matched noise voxel sample by normalizing the stimulus voxel matrix thus unmasking a highly discriminative neural profile per stimulus. Projection of this new voxel space (ROI) to a smaller set of dimensions (with e.g., non-metric Multidimensional scaling), the relational information takes a unique geometric form revealing functional relationships between sets of stimuli, defined by R. Shepard, as *second-order isomorphisms (SOI)*. In the case of colors the SOI appears as a nearly equally spaced set of wavelengths arranged in a color wheel, with a gap between the “purples” and “reds” (consistent with the gap in the original Ekman’s color set). In the case of letters, a cluster space resulted from the decorrelated voxel neural profiles, which matched the phrase structure of the mnemonic used for more than 100 years to teach children the alphabet (across multiple languages), *The Alphabet Song*.

## Introduction

One of the earliest and enduring questions for those interested in human cognition is how the external world is internalized, i.e., how the brain maps objective information to subjective experience. Neuroimaging techniques such as fMRI (functional magnetic resonance imaging), provide the opportunity to answer this question. The typical answer from neuroimaging studies has been that some brain regions are functionally dedicated to processing certain kinds of information. Examples of such regions are the fusiform “face area” or the extrastriate “place area” as well regions activated by shape (*1*), objects (*2*), peripersonal space and body ownership (*3*), and words (*4*).

However, the claim that certain brain regions uniquely process specific kinds of knowledge (e.g., faces) has become controversial, and there is mounting evidence that knowledge about the world is represented in the brain in a distributed form (as networks) rather than in localized brain regions (*5*, *6*) consistent with recent brain connectivity research (*6*, *7*). Specifically, there is a general consensus that an “object” is represented in the brain, not as a complete entity (e.g., a “house”, a “face”, etc), but rather as a pattern of activation distributed over neurons and therefore voxels (3D brain units). Thus, a set of voxels can contribute to the representation of multiple objects in much the same way that a set of musical notes can contribute to multiple chords, or an alphabet can contribute to multiple words (*6*).

Researchers generally agree that the representation of objects in the brain does not appear as a first order isomorphism (FOI) as that proposed by Gestalt psychologists (*8*) who argued that brain representation should share physical resemblance with the object being perceived, either concretely (physical properties) or abstractly (feature configuration). There are, of course many examples of direct sensory mapping to brain topography (e.g., retinotopy and tonotopy), however, these are mappings of physical sensory features (amplitude or spectral properties) rather than perceptual entities (objects). This type of first order mapping is important for understanding sensory processing, but it does not capture the subjective experience dependent on a feature configuration or relations inherent to perceived objects.

### Second Order Isomorphism (SOI)

There is an alternative to the Gestalt view of a physical resemblance between the external world and the representation of physical objects in the brain (*9*, *10*). In this account, brain representation is a second order isomorphism (SOI) in which the relation of an object to other objects, rather than the physical properties of that object, provides the basis for representing the physical world in the brain:

> “*…the <second order> isomorphism should be sought not so much between any individual stimulus and its corresponding internal representation but between the <u>set</u> of potential stimuli and the <u>set</u> of their corresponding representations. It is, I think, a kind of metric of similarity among stimuli that is largely preserved in the functional relations among inner constructions.”* (9; p 288).

There is strong support from behavioral studies (subject performance in tasks) that second order isomorphism serves a functional, relational organization for perception, motor control, and cognition (*11*–*13*). In a classic demonstration, Shepard (*12*) applied nm-MDS (*14*, *15*), which is neither equivalent nor statistically related to PCA or ICA, to similarity judgements of colors collected in another study (*16*). The nm-MDS analysis produced a 2D representation that recovered the relation of colors represented by the color wheel (*17*), including the physical gap between the smallest wavelength and the largest in the stimulus set. The 2D embedding of the color objects are functional in that color relations could be “read” directly from the structure itself.

Theoretically, Shepard’s approach could be used to extract a second order isomorphism directly from brain activation data provided a neural distance matrix could be extracted from fMRI data as is done with behavioral similarity judgements. However, detecting SOIs using brain activation (fMRI) poses many more challenges than extracting SOIs from behavioral data. One of the most critical challenges involves the enormous noise that exists in neuroimaging data. For example, fMRI data typically reflects a very small signal to noise ratio; generally only 4-6% of BOLD (blood oxygenation level dependency) activation is associated with a typical stimulus presentation (*18*, *19*). Much of the noise in which the signal is embedded (e.g., diffusion in space and time of the hemodynamic response to a stimulus) affects autocorrelation within a voxel time series, and of interest here, stimulus *cross-correlation*, making it nearly impossible to decode individually presented stimuli or their relations. The new method we describe here, provides significant decorrelation at the stimulus level, heretofore unseen.

### Cross-Correlation and Distributed Encoding

In a typical fMRI study, the GLM (General Linear Model) is used to model the voxel activation time series for a given stimulus class against the temporal presentation of individual exemplars of that stimulus class. Voxels that are recruited when processing a given stimulus class will respond more than those not recruited, thereby providing information about which brain region is involved during a given condition. In general, the goal of these kind of analyses is to differentiate one experimental condition (e.g., cat exemplars) from another experimental condition (e.g., dog exemplars) in voxel time series data collected throughout the brain. Because it is generally not of interest to distinguish individual exemplars of the presented stimulus class (but see 20), a primary concern is the autocorrelation of voxels across conditions; a reason why many experimental designs incorporate relatively long inter-block intervals or staggered event related stimulus presentations that can be phase aligned for averaging. Standard decorrelation methods (PCA, ICA, etc.) are often used to decorrelate activation from individual conditions (with a category of stimuli), forcing the off-diagonal values towards zero and effectively masking the possible relations within individual conditions of the presented stimuli themselves and are incapable in unmasking the stimulus relations that this boosting method can accomplish.

Because our interest was in extracting the relation among individual stimuli, rather than among stimulus classes, standard denoising techniques could not be used. Instead, we developed a model in which it is assumed that a distributed representation of neural features (V1, V4 or extrastriate for example; *6, 21,* exists generally) in the brain. The model is designed to reduce the noise in cross-correlation and therefore *boost* each individual stimulus presentation while not necessarily affecting the autocorrelation of each voxel time series (see supplemental section for more detail on decorrelation).

Figure 1 displays the model workflow used to recover the second order isomorphic representation of colors. This workflow is identical to that used for letter processing with the exception of the brain region from which the voxel time series data are selected. The first step of the workflow is extracting a stimulus by voxel matrix using brain activation (voxel time series data). From this matrix, a voxel level gaussian kernel smoother is created over stimulus voxel profiles. These profiles are further aggregated by averaging over all individual stimulus profiles creating a column marginal. (Although the normalization works with the smoothed data matrix, further noise reduction at the voxel level is often even more effective).

**Figure 1.**
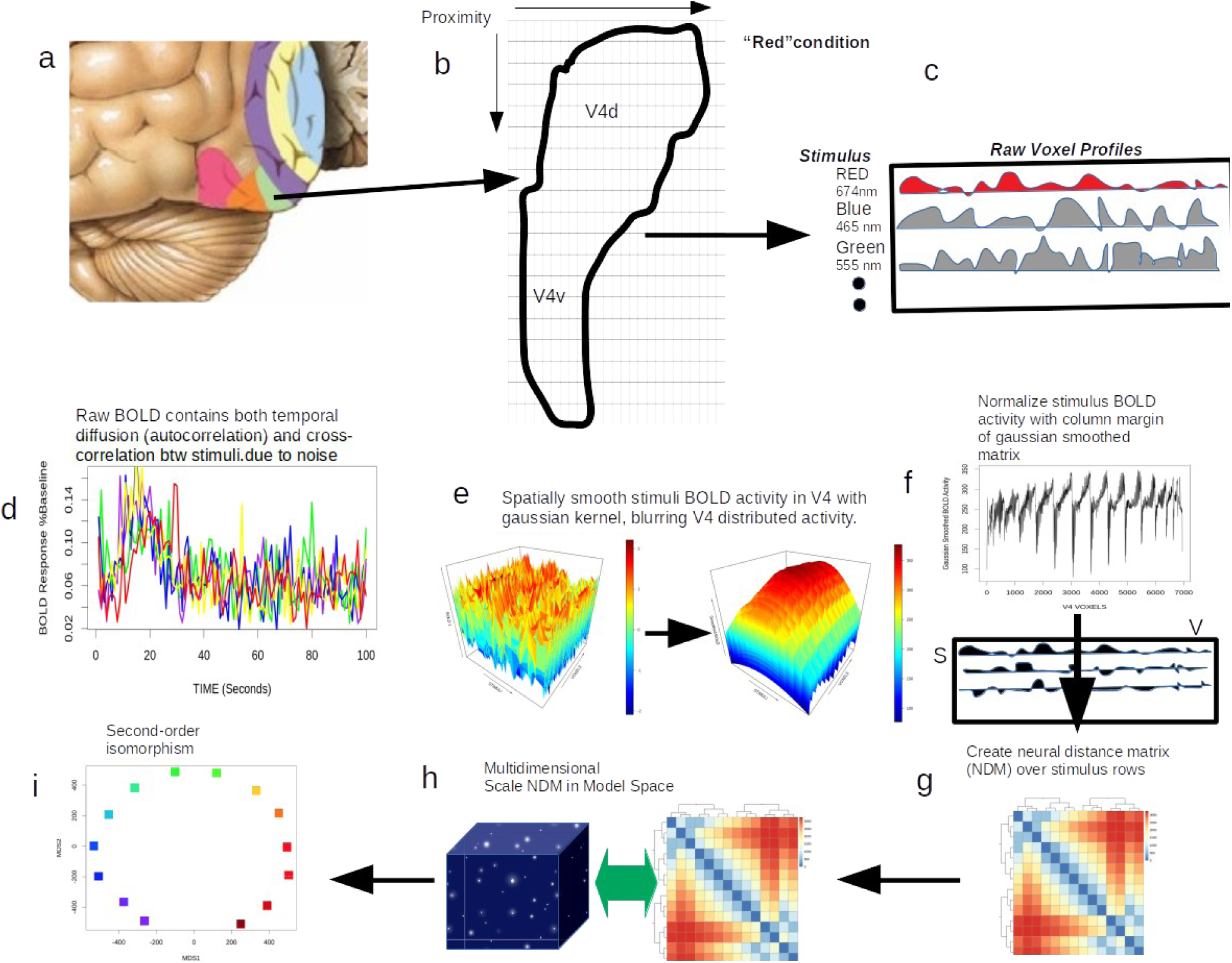
We show the model workflow of decoding various cortical areas and construction the SOI. Standard tools for tissue extraction tools (*22*) are used with a probabilistic mask (Juelich masks; see supplemental material) to extract a regular grid of voxels during a specific condition. In the first two panels (a,b; upper left corner) we show area V4 overlayed with a regular grid in which fsl-meants has extracted ∼6000 voxels from this particular V4 mask. Repeating this extraction for each individual stimulus produces a stimulus condition (for colors 14, for letters 25) by voxel matrix shown in the top left corner (c). In this case each stimulus condition row, which in this example is a single wavelength (CIE) presented to the subject for 6 seconds (see supplemental for more detail). In panel d, we show the typical hemodynamic response with additive noise per color condition. Note the enormous overlap between colors and their cross-correlation shown in figure 2d. In panel e, we show the gaussian kernel smoother over the stimulus matrix columns blurring the voxel profiles per stimulus condition.This is further aggregated as a margin average, again over voxels shown in panel d and then used to normalize the original stimulus matrix (c). At this point the stimulus matrix is used to calculate the neural distance matrix (g). Finally in panel h, this matrix is submitted to nm-MDS producing a solution with low stress (high variance accounted for) in 2 dimensions.Plotting the solution reveals the color second-order isomorhpism (panel i).

This marginal is then used to normalize the stimulus profiles over stimuli and to recycle (**n stimulus times**) the smaller smoothed vector over the larger original raw stimulus data matrix. This produces a 40-60% stimulus decorrelation (signal boosting) from the base values (see Figure 2) without forcing a complete orthogonal decorrelation between stimuli. From the newly decorrelated stimulus matrix, a row-by row neural (voxel) distance matrix (NDM) can be computed using Euclidean distance. That NDM is submitted to nm-MDS analysis which results in the n-dimensional SOI.

**Figure 2.**
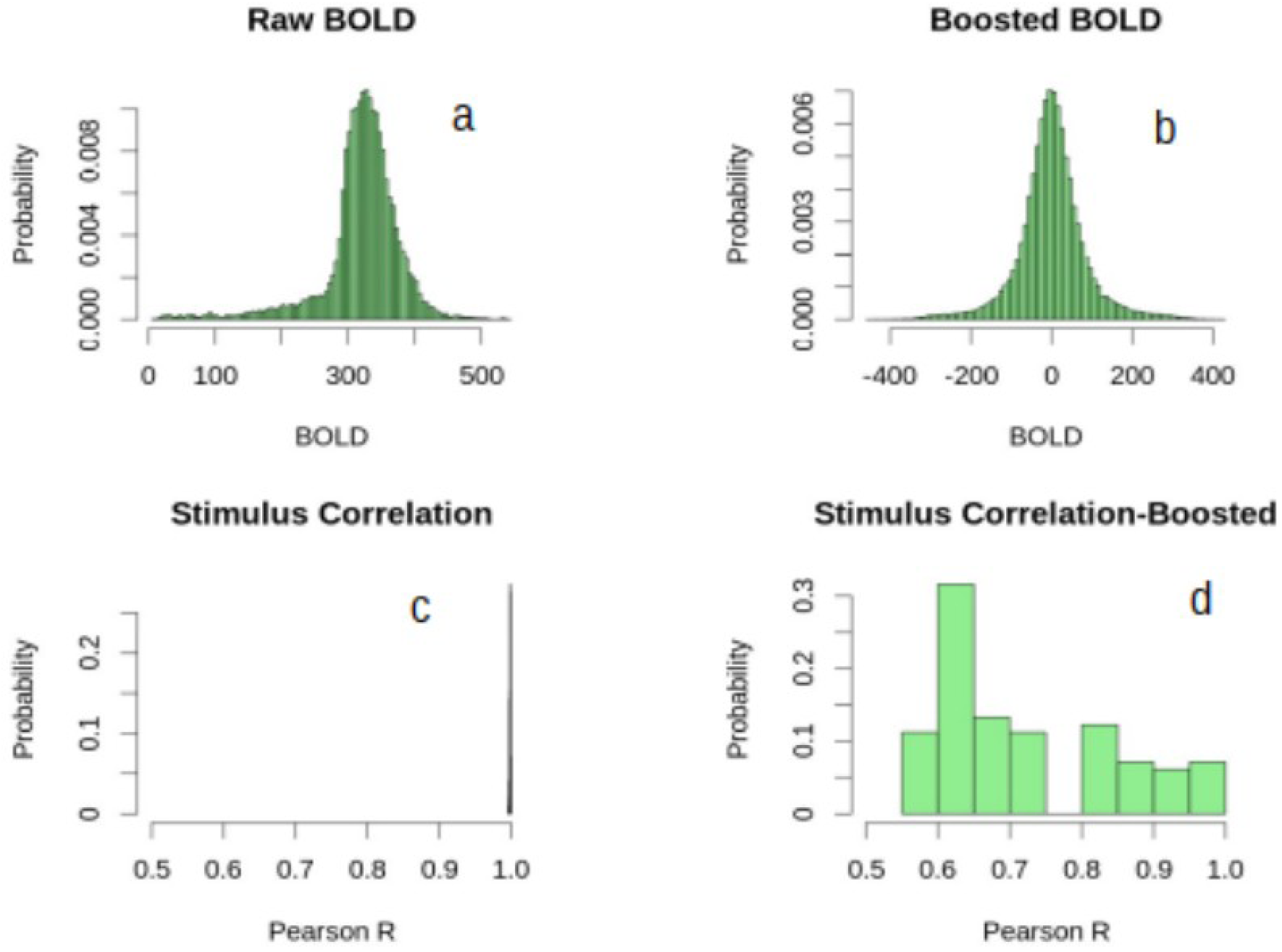
Boosting results for fMRI. In this panel we show the results of the boosting model on fMRI from the single color presentation stimuli. In this case we start with a standard BOLD distribution (Figure 1a) which shows a classic long tail with left skew and a peak of noise just prior to the signal bump starting around 450 on the x axis. Once the boosting model is applied to the raw BOLD, the distribution appears to normalize and is scaled in mean and variance (Figure 1b). The original raw BOLD correlation of the stimulus x voxel matrix is shown in Figure 1c, which indicates the effect of BOLD noise on the cross correlation of the stimuli over voxels with a range of .997-1.0. Once boosted there is a sphering effect on the stimulus correlations resulting in a typical range of .5-1.0 (also see supplemental-S4).

Mathematically, we can describe the model in the following equations starting with the kernel gaussian smoother over the voxels patterns by stimulus condition matrix.:

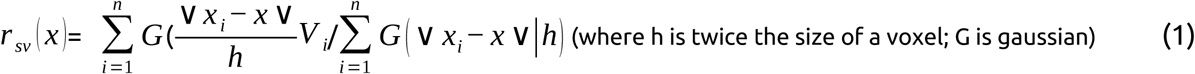

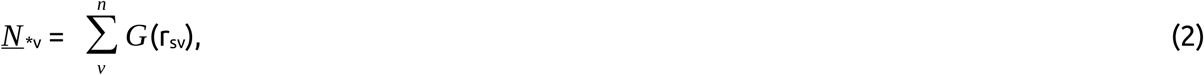

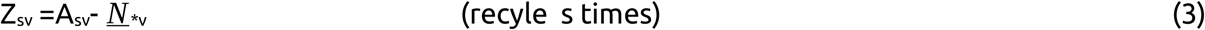

This filtering model was applied to two types of physical stimuli (colors and letters) which were chosen because subjective representations had been extracted for each type of stimulus in earlier research using similarity judgement data (*12*, *23*). In these earlier studies, subjective representation of colors mirrored that of the color wheel following nm-MDS analysis and similarity judgements between pairs of individual letters produced clusters representing different physical/spatial features of the letters (round, tail, branch, arched, vertical) after submission to clustering analysis. We replicated these results for colors and letters In independent studies using behavioral similarity judgements alone as was done in the original studies (supplementary material S1abcd).

Given the nature of the SOI representations extracted from behavioral data, it begs the question of what form the SOI extracted from neural data might take? Are the neural SOIs likely to have higher fidelity or some sort of cannoical geometries? In the case of colors, we extracted a neural distance matrix (NDM) and submitted it to nm-MDS. We found an SOI that was highly consistent with the color wheel, and remarkably, one which more robustly mimicked the physical relation among color stimuli than did that arising in the original study or our behavioral replication (see replication in Supplemental material). Shown in Figure 3 are the SOIs based on fMRI data obtained from four different subjects. Notably, the SOIs are highly consistent with the color wheel and vary minimally across subjects.

**Figure 3.**
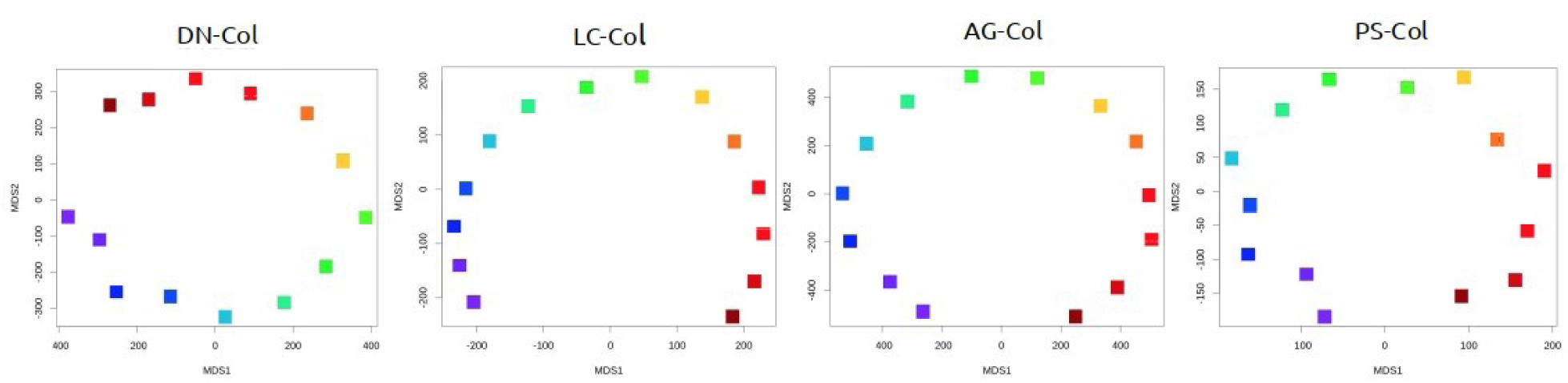
Color geometry SOIs recovered from nMDS after boosting stimulus signals. These color circles represent the response of four individual subjects to single color presentation as described in the procedures section. Compared to their individual behavioral results, which are averaged over 11 subjects, the increase in fidelity from the direct neural distance matrix is dramatic. Where the averaging of behavioral data is required to begin to observe the convex ordered “color wheel”, a single subject shows from cortical response only a clearly defined nearly closed circle with the original Ekman stimulus gap in the 14 color wavelengths chosen, while the rest of the pairs of stimuli are nearly equal spacing.

In the letter case, we also found high consistency among subjects in the SOIs obtained. In this case, we used hierarchical cluster analysis in order to replicate the original study (23). However, to our surprise the letter SOI obtained from the brain activation data was not based on visual features of the letters as seen in the original study or in our replication using similarity judgements.Rather, the cluster dendrograms (Figure 4)match (cutting the dendrogram at k=5, and crosstabs with alphabet song—100%,99%,97%,98%) the phrase structure used in the “Alphabet Song” (Figure 5), a mnemonic universally used to learn the English alphabet. Clustering solutions obtained for the four subjects differed minimally, as the color coding in Figure 4 shows.

In fact, Errors in the color coding appear to occur at the transitions in musical phrase structure of the Alphabet Song (Figure 5), such as “abcdefg”, “hijkl”, where “g” or “l” might appear in an earlier or later sequential set, suggesting that a continuous geometric solution might better fit the letter NDM. Submitting the letter NDM to nm-MDS produced solutions from two to seven dimensions (based on “Shepard plots”). A three dimensional nm-MDS solution for the same subjects (Figure 6) reveals a complex sequential structure in which points of inflection in the data coincide with the phrase boundaries (not necessarily order) of the Alphabet Song, similar to that found with the cluster analysis.

**Figure 4.**
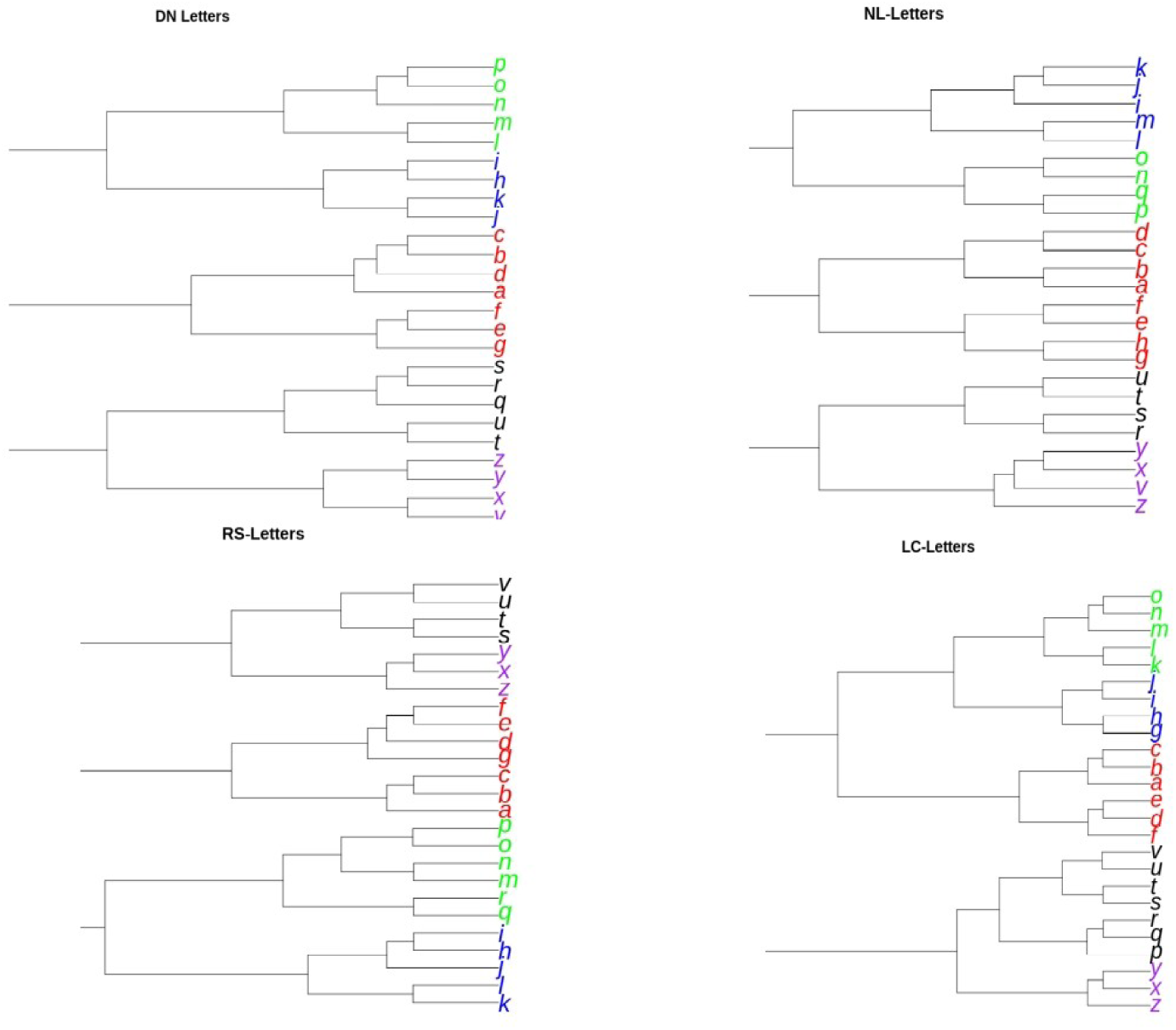
Letter cluster SOIs recovered from hierarchical cluster analysis (euclidean distance, Wards grouping rule) after boosting stimulus signals. The cluster dendrograms are fit to each individual Neural Distance matrix producing 5 clear cluster groups that per group tend to be sequential alphabetically, but more surprising, they follow the phrase structure or prosody of the the “Alphabet Song” (in Figure 4) which is a universally known mnemonic for learning the english (latin) alphabet. The phrase structure is grouped as the RED group (“A”,“B”,“C”,“D”, “E”,“F”,“G”), then the BLUE GROUP (“H”,“I”,“J”,“K”), then the GREEN group (“L”,“M”,“N”, “O”,“P”), BLACK group (“Q”,“R”,“S” & “T”, “U”) and finally the PURPLE group (“V”, “X”, “Y”, “Z”). Except for “W”, which was excluded from the original sweedish study (as “W” is rare in sweedish), we see a close match between the Alphabet song and the sequential groupings from the hierarchical cluster analysis. Errors in the mapping tend to only occur at the boundaries of the phrase structure (see Figure 4). Another interesting result is that the behavioral similarity judgments also submitted to hierarchical cluster analysis (euclidean distance and Wards grouping rule replicated the 1969 study producing 5 cluster groups based on visual similarity. This implies a dissociation from behavioral SOIs and neural SOIs.

**Figure 5.**
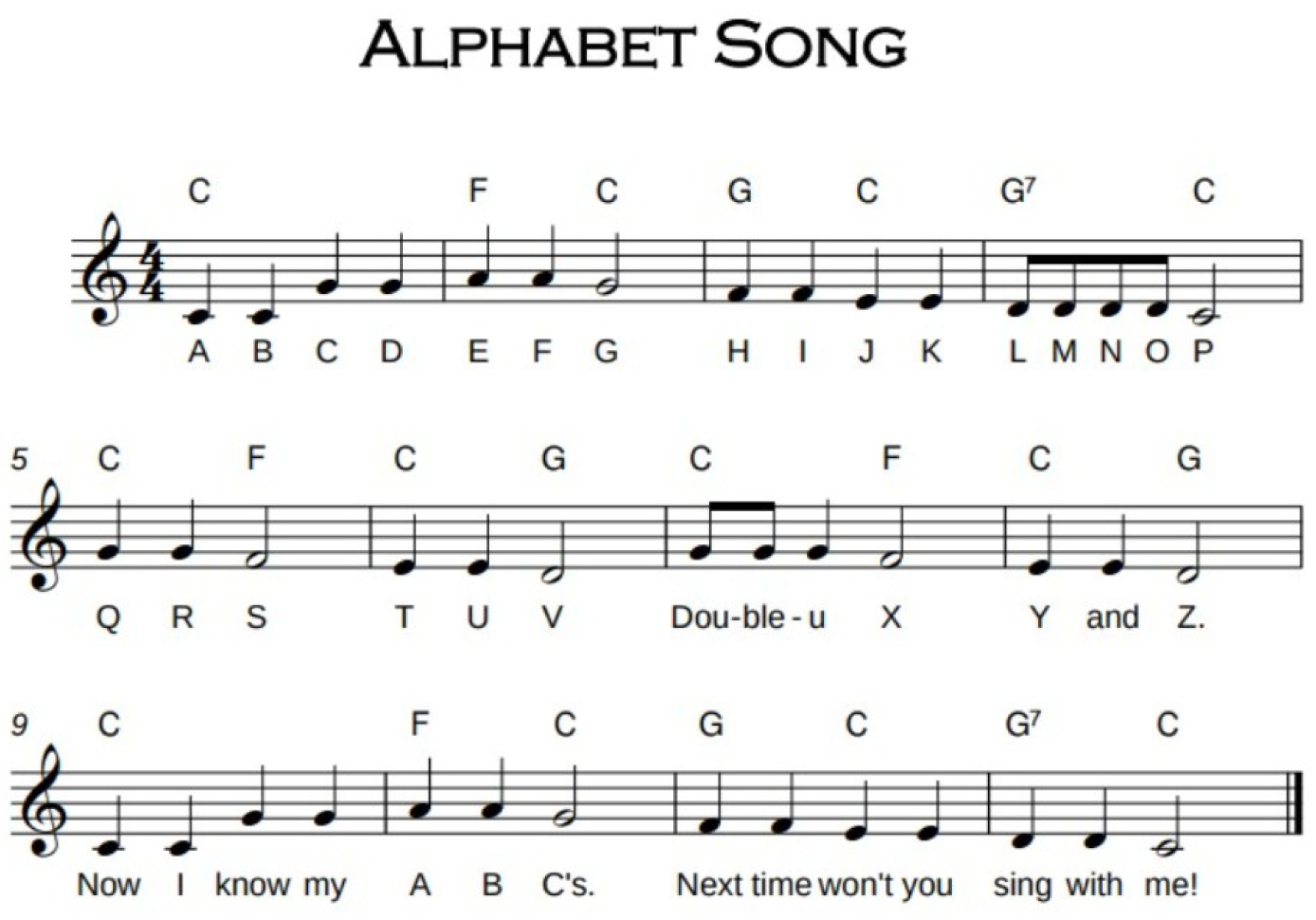
“The Alphabet Song” which is set to the melody of “twinkle, twinkle little star”, has provided a nearly universal way of teaching children letters for more than 100 years. The phrase structure, or prosody of the song follows the following groups (melody rising) A, B, C, D, E, F G <pause> H, I, J, K, <pause> L, M, N, O, P <pause> Q, R. S, and T, U, V <pause> double-U, X, Y, Z. Subjects all debriefed after the experiment (once we identified the SOI) were asked had they heard of the Alphabet song or could they sing it. Every subject immediately began singing with the same phrase structure shown in this prosody/melody.

**Figure 6.**
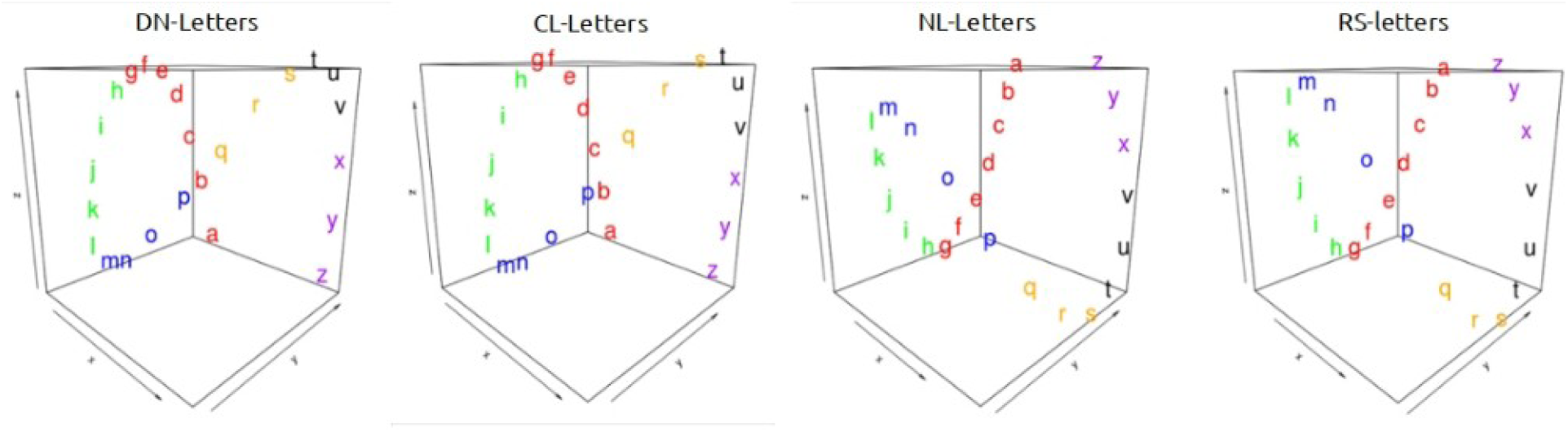
Because the NDM in the hierarchical cluster dendrogram appeared to produce errors near each boundary of the phrase structure in Figure 4, it seemed likely that the SOI may be more continuous or possibly higher dimensional. We therefore submitted the same NDM to nMDS, which produced a complex higher dimensional structure with more than 2 dimensions and less than 7 (given the stress scree plot). The SOI appears in all four subjects as a “figure 8” showing that the Alphabet Song phrase structure is coded by direction change in the phase space (color coded for the Alphabet Song) or inflection point more generally. Possibly the SOI codes letter frequencies or higher order lexcial structure in words.

## Discussion

Whereas first order isomorphism can exist for sensory information (e.g., energy associated with light or sound), the representation of entities with holistic feature integration (e.g., objects) requires a higher order isomorphism that can capture the relation among objects in the external world and potential behavior. Most physical objects can not be categorized by a set of necessary and sufficient features, but rather are defined contextually in comparison with other objects and are ill-defined, organized by typicality. Cats and dogs are more similar than are cats and cows, yet all are breathing, four-legged, terrestrial, non-flying, domesticated mammals. Representations of second order isomorphism must also provide leverage in cognitive/perceptual processing, decision processing, feature judgement and motor schemata.

The existence of cognitive representation based on second order isomorphism has been readily established using behavioral responses (similarity judgements). However, extracting second order isomorphism from brain activation provides a much greater challenge. One recent attempt of color decoding from fMRI data used MVPA, PCA, and a filter bank of six equally spaced ordered color channels to decode a color wheel from V4 activation (*24*). We too were able to find various decodings of the color wheel without applying the boosting model described in this report. However, when the boosting model is not used to initially decorrelate fMRI data, the results in nm-MDS or a PCA space can often be convex but do not generally preserve the relation of hues in the color wheel or its circular form.

Using the boosting model, we have demonstrated how an SOI can be revealed for two very different types of stimuli: colors and letters of the alphabet. In the case of colors, the SOI obtained with brain activation data completely replicated that obtained with behavioral data. In the case of letters, which have a higher dimensional manifold, we found an SOI that mirrors the “Alphabet Song” even though the SOI derived from similarity judgements obtained from the same subjects replicated behavioral studies showing visual letter shape similarity cluster outcome.

This dissociation in the letter SOIs obtained from behavioral and fMRI data may arise because letters naturally have a higher dimensional manifold than do colors (or they are learned), leading to more possible configurations in the way they can be represented. When subjects are asked to judge the similarity of visually presented stimuli, visual features are the most salient criteria to use and in fact result in SOIs such as those revealed in the behavioral letter studies. However, visual features are only one dimension on which letter similarity can be judged. Phonology, for example, is another dimension on which letter similarity could be judged. The SOI derived from the fMRI data was a complex representation that incorporates several dimensions of letter stimuli and suggests that the brain constructs a representation of the physical world that reflects the various ways in which sets of stimuli contrain one another as dictated by the kind of experience acquired, thus creating a second-order isomorphism between external stimuli and their internal represenation.

## Supporting information

Decoding_supplemental

