## Supplementary material for "DECODING SECOND ORDER ISOMORPHISMS IN THE BRAIN: The case of colors and letters": Decoding_supplemental

### Supplemental Material: Decoding Second Order Isomorphisms in the Brain

#### Methods

***Color experiment:***

*Subjects.*

A total of 15 (mean age = 26.3, female = 6) participants were recruited to participate in the behavioral replication and 4 participants (mean age = 26; female = 2) for an fMRI version of the original Ekman color task. Participants were prescreened for possible colorblindness using the Ishihara color test (Ishihara, 1917) and gave informed consent in accordance with the Rutgers University Institutional Review Board for the Protection of Human Subjects in Research and either volunteered or received course credit for their participation.

*Stimuli.* Square color stimuli (31x31 pixels) were created based on the wavelengths of the original 14 colors used in the Ekman (1956) study. The fourteen wavelengths were 434nm, 445nm, 465nm, 472nm, 490nm, 504nm, 537nm, 555nm, 584nm, 600nm, 610nm, 628nm, 651nm, 674nm. We converted the color wavelength values used by Ekman to the tri-stimulus color space CIE XYZ to create digital color stimuli that most closely resemble the perception of colors by the human eye (CIE, 1931) . The stimuli were displayed on a 12-inch retina display MacBook (2304 x 1440 resolution at 226 pixels per inch). These procedures replicated behavioral similarity judgments of the Ekman (1954) study (see figure S1).

*Task.* To assure that participants were familiar and consistent enough with using the similarity scale, they received 2-3 training runs of the task prior to performing it in the fMRI scanner. The experimental task was presented using PsychoPy to participants in the MR scanner. The task consisted of two parts, the first to collect the neural data and the second for the behavioral data. In the first part, participants saw two runs of each color stimulus individually (2x14 colors) in a block design. Each trial consisted of first seeing either a cross or a hash (coupled with an ISI of 3-4s) and then a color stimulus (6s). As an attention check, participants were instructed to indicate via button press whether they saw a cross or a hash. The order of both the colors and the cross/hash presentations were randomized per participant and repeated over two separate runs.

In the second part, participants were instructed to rate the visual similarity between all possible color pairs, including the diagonal of identical pairs (105 pairs total; (14*13)/2 + 14, including the diagonal of identical pairs). A fixation cross was presented with the ISI (1-2s), followed by the presentation of colors pairs (4s) and a similarity scale that appeared after the first two seconds (2s; 1-5 with 1=no similarity; 5=identical). The order of pairs was randomized for each participant.

#### *Letter experiment:*

*Subjects.* Four participants (mean age=28; female=2) were recruited for the fMRI experiment of the letters, replicating the behavioral results of Kuennapas and Janson (1969). All subjects were prescreened for no history of dyslexia or reading impairments.

*Stimuli*. We used the same set of Latin alphabet letters as in the Kuennapas and Janson (1969) study, with the exception of three Swedish (å, ä, ö) letters that do not exist in the English alphabet as well as "w" due to its rare use in Sweedish. The final set of letters (25 letters) included all remaining letters in the English alphabet. Subjects were asked to rate the visual form similarity of the pairs of letters. These procedures replicated the original Kuennapas and Janson (1969) based on the shape of the visual features of letters (S3).

*Task.* The experimental task was presented using PsychoPy. Participants were instructed to rate the visual similarity between all possible letter pairs, including the diagonal of identical pairs (325 pairs total). Identical to the design of the color task, a fixation cross was coupled with the ISI (1-2s), followed by the presentation of letter pairs (4s) and a similarity scale that appeared after the first two seconds (2s; 1-10 with 1=no similarity; 10=identical). The order of pairs was randomized for each participant. Due to the 4 fold increase in judgments compared to the color experiment and the likely increase in number of dimensions (which could also be auditory as opposed to visual) we were concerned about the potential attention variation over two replications of single letter presentation and pilot studies seemed to indicate that priming the single letter presentations with the similarity task would improve signal to noise. Consequently, the letter task in the scanner presented single letters chosen from the pairs of letters in the similarity judgment task in which the subject indicated whether the single letter presentation was on the “left” or the “right” of the subsequent pair that was presented.

**fMRI data acquisition parameters**

All neuroimaging data was collected at the Rutgers University Brain Imaging Center with a Siemens TRIO 3T scanner and 32 channel coil. The experimental tasks were presented using PsychoPy, where stimuli were projected on a screen at the back of the bore which was visible through a mirror mounted on the head coil. Participants’ responses were collected via presses on a fMRI compatible button box. We collected anatomical MRI data (T1-weighted 176 1mm slices) as well as functional data (Multi-Band), based on 35 contiguous slices (1 mm voxel resolution) using a Multi-Band sequence (TR = 835 ms). Preprocessing, spatial normalization to standard MNI space and analysis of fMRI data was conducted in FSL (Smith et al., 2004) and R (R Development Core Team, 2008).

**ROIs** For the color data, anatomically defined masks of the region of interest (ROIs) V4 and the control ROI Auditory Cortex (thresholded at 60% probability) were created using the Juelich Histological atlas. The ROI for the letters was defined as the Extrastriate cortex (V2-V5) and primary somatosensory cortex was used as a control ROI. Both ROIs were anatomically defined based on the Juelich histological atlas and thresholded at 50% probability (see masks below S10-S14).

**Behavioral Results.**

*Behavioral Replication of Colors and Letters.*

Behaviorally, we replicated the similarity judgements from 15 (mean age = 26.3 , female = 6) subjects asked to provide a similarity judgement (1-5) on 105 pairs of 14 Ekman color stimuli (Ekman 1954), subsequently scaled by Shepard (1962), using nm-MDS. The average of subjects is shown in Figure S1 (ab). Panel S1a shows the resultant color wheel of the behavioral similarity judgements matching the original Ekman-Shepard nm-MDS, while figure S1b shows the best fit in the 2D nMDS space ($r^2$=.96) Figure S1 shows the same type of results per group average

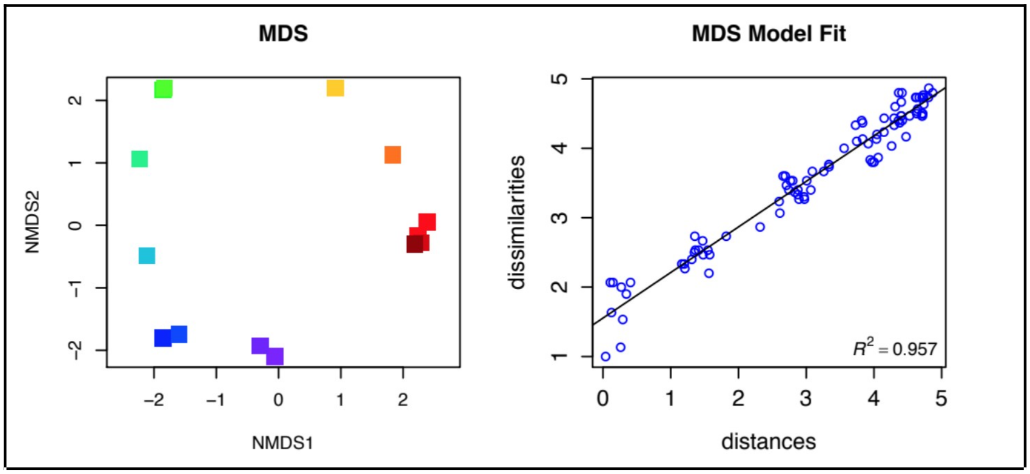


Figure S1. Color behavioral replication of Ekman/Shepard color wheel. On the left are 15 subjects similarity judgments which were averaged across distance matrices and then submitted to a nMDS, producing a color wheel, with some clustering in very similar wavelengths. The Shepard plot (right) showed that the recovered distances in the model nMDS fit closely followed the subject similarity (or rather dissimilarity) judgments.

for the behavioral color data collected during fMRI scanning. Figure S2 shows the same type of work-flow of the letter similarity group average based on visual similarity judgements (1-10) of 325 letter pairs collected during fMRI scanning replicating the Kuennapas and

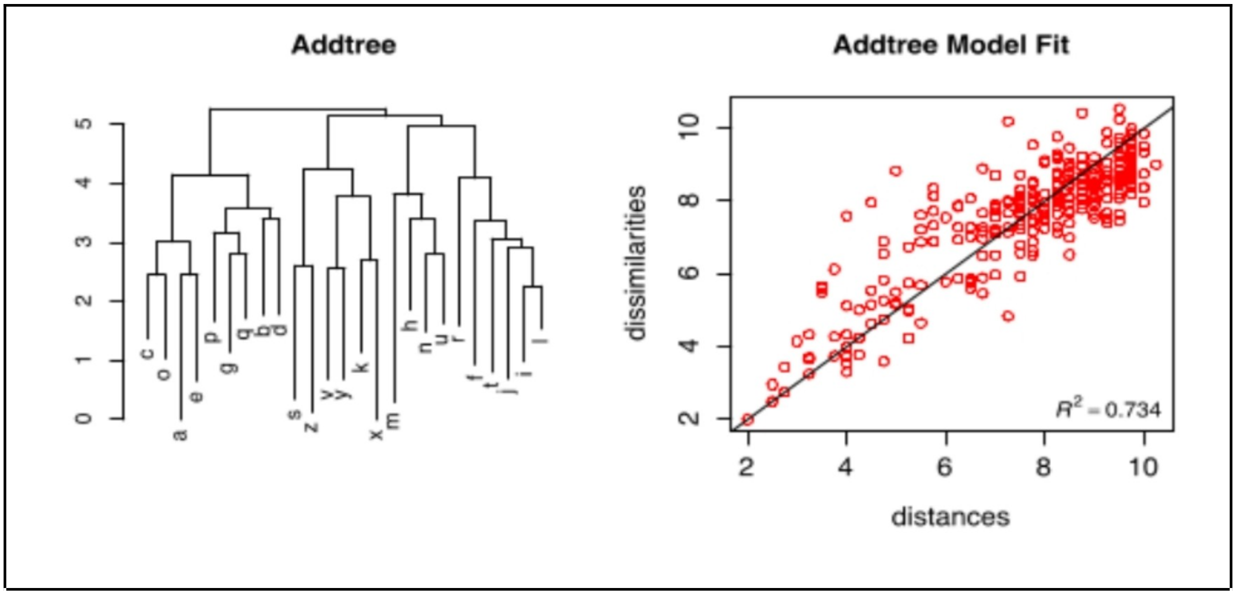


Figure S2. Letter behavioral replications for Letters when subjects are asked to produce similarity judgments on visual shape similarity. The resultant cluster dendrogram (left) due to ADDTREE, are five different clusters including from left to right—*round, tailed, branch, arched, vertical*), and a Shepard plot (riget) showing a fit of 73%, higher then the best fitting nm-MDS.

Janson (1969) results, showing 5 major visual feature clusters *(left to right: round, tail, branch, arched, vertical)* and cutting the dendrogram at h=4.5 with an Addtree fit of $r^2$ =.73).

***Colors from fMRI.*** All experimental procedures were repeated in the scanner with exactly the same subject instructions (including the visual shape instruction for the letter stimuli). Subjects in the color task were asked to do the same similarity judgments and then a series of two replications of the 14 colors (i.e., 28 total) shown in random order after a fixation. A neural distance matrix was calculated from the extracted V4 (6950 voxels; V4 mask) stimulus by voxel after decorrelation. This neural distance matrix was submitted to nMDS with resultant 2D scaling shown in the main report, Figure 2 (a,b,c,d) for all 4 subjects. Note the recovery of the color wheel per subject in original Ekman (1956) stimulus order from voxel activity alone.

***Letters from fMRI.*** For the letters as with colors, all procedures in the scanner were the same as those in behavioral experiments. The BOLD acquisition was also identical to the colors fMRI and a matrix of 25 letters by 8624 voxels from extrastriate areas including (V2-V5 Figure 4). In Figure 4 (a, b,c,d) are shown the resultant cluster analysis of letters which show a *sequential* as opposed visual shape similarity as in the behavioral replications. In closer examination, there is also a musical phrase structure based on the cluster solution. In this case, we have color coded (5 cluster solution) the phrases from the "Alphabet Song", a universal (in most languages) mnemonic for learning the sequence of the alphabet, the color groups (*red, green, black, purple, orange*) indicate an almost perfect match to the melody phrase structuring of the song (shown in the main report, Figure 4).

***Decoding/Decorrelation methods.*** Generally, a decorrelation process is defined to either reduce the autocorrelation within a signal or the cross-correlation between signals. A common approach to decorrelation due to autocorrelation is to create a whitening matrix based on the covariance (voxels time series) of the original sample and/or further extracting the eigenvectors of the covariance matrix to filter the original data matrix. Raw BOLD from fMRI measures are known to be autocorrelated over time due to the hemodynamic diffusion over conditions or stimulus presentations. Another more difficult association is cross-correlation, again due to the hemodynamic diffusion which in this case comprises the "background" noise, which every condition or stimulus presentation is sharing, and consequently increasing their cross-correlation. On the other hand, Cross-correlation (of Voxel time series) have proven to be important in Brain-Connectivity, creating a functional correlation between voxels or clusters of voxels ( ROIs). Since the signal activity relative to the BOLD can be a small as 2% of the stimulus/condition, the cross-correlation is generally pernicious and ignored in standard neuroimaging protocols (FSL,AFNI, MRI-prep etc). In fact, the larger focus of decorrelation in neuroimaging has been in service for the detection of the brain location of a given stimulus *class* presentation. A productive tactic has been to create a statistical voxel contrast between two conditions that share a minimal and/or diagnostic overlapping feature structure. This strategy, unfortunately can leave the stimulus conditions cross-correlated due to the overwhelming shared noise background (see Figure S3a). This has not at the same time been a problem for the neuroimaging community which has Standard whitening approaches using the eigenvectors from the covariance matrix will produce a reduction in cross-correlation causing the transformed covariance matrix to tend towards *zero* off-diagonal entries. At the same time, unfortunately, this kind of decorrelation can be very aggressive leaving the stimuli orthogonal (i.e. PCA see Figure S3b ) or more strongly statistically independent (i.e. ICA) very likely disrupting any underlying SOI. The usual model assumes that data matrix X is a linear function, L, of signal and additive noise:

$$X = L(S + N)$$

assuming that that X is has a non-singular covariance matrix $\Sigma$ (which may be estimated with maximum likelihood shown figure S3c) with centered mean 0, then a transformation:

$Z = WX, where\ W = \Sigma^{(-1/2)}$ will result in Z to be a whitened random matrix with unit diagonal covariance. This sort of whitening process is often applied to voxels as a preprocessing state, in order to increase the brain location detection and signal sharpening.

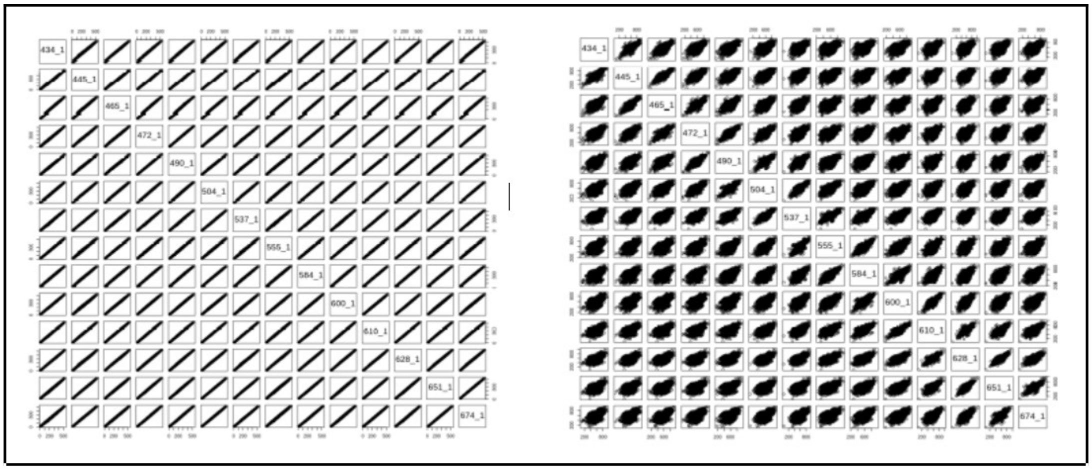

Figure S4. In this figure we show the dramatic sphering effect from the original wavelengths in the diagonal (left) showing high off-diagonal correlation (due to fMRI noise) and their subsequent decorrelation (right) due to the boosting algorithm.

In the present decorrelation model, we focus on the whitening across voxels per stimuli. The row voxels over stimuli represent a profile of brain activity in the regional map where time series are extracted. This was, in fact, the rationale for MVPA (Haxby, et al, 2001; Hanson et al, 2009), where voxel patterns are used to detect and classify regional brain areas as stimulus predictive conditional on the MVPA model. Here again, the focus is on separating stimulus locations ("face areas" etc..) from other stimulus locations ("place areas" etc..) rather then identifying relational structure between stimuli. The insight in the present model, is that the profile across voxels within stimuli rows can be noise "scrubbed" to only lessen the correlation between patterns and boost the specific stimulus signals (resulting in a "sphering" see Figure S4a and S4b). We have found this approach to be most effective, with the original raw voxel matrix, rather than a reduced PCA or otherwise projected set of voxel components, because methods like PCA are linear and/or highly aggressive in reduction into a smaller set of components. Choosing the voxels with the largest activity can also reduce the number of voxels in the profile that are decorrelated (also note that removing too many voxels will destroy the SOI) . However, in the present application we used the same number of voxels for every subject, in order to ensure a common basis for establishing the efficacy of the algorithm and to avoid any accidental bias.

The decorrelation algorithm creates a mean vector from each voxel, resulting in an aggregate of all stimuli per row. So, for example, in terms of the colors, a mean vector of all 14 (or 28 presentations) colors expressed as voxel patterns in the marginal is subtracted (or added-any noise perturbation is effective) to each of 14 rows, with a much larger ~6000 mean vector, which is consequently recycled over the columns with each 14 of the rows. This has the effect of whitening the row vectors and decorrelating the individual stimuli as shown in the main report in Figures 2, 3, and 5. We should note that not any kind of noise will work, for example, creating a similarly sized gaussian matrix and deriving an aggregate vector, resulted in the stimulus rows unchanged in their cross-correlation. We also tried to use mean noise samples between subjects. Although, various subject's noise vectors could be used decorrelate other subjects , this approach was not consistent with all pairs of subjects causing the decorrelation to fail arbitrarily, suggesting some specificity in matched noise samples either subject or session dependent.

***Controls: Other brain areas***

***(1) Colors***

We examined other brain areas to control for cortical sensitivity location. We focus on cortical areas that are unlikely to produce a visual response such as auditory cortex. We show , for example, in S5a that nMDS of auditory cortical areas (Herchel's gyrus mask) after decorrelation (looked similar prior to decorrelation. Next we show nm-MDS without boosting: First in Figure S5b if the NDM is not pre-processed with the boosting method the scaling outcome recovers neither the convex color 2D structure nor the order of the wave-lengths, which is consistent across all subjects.

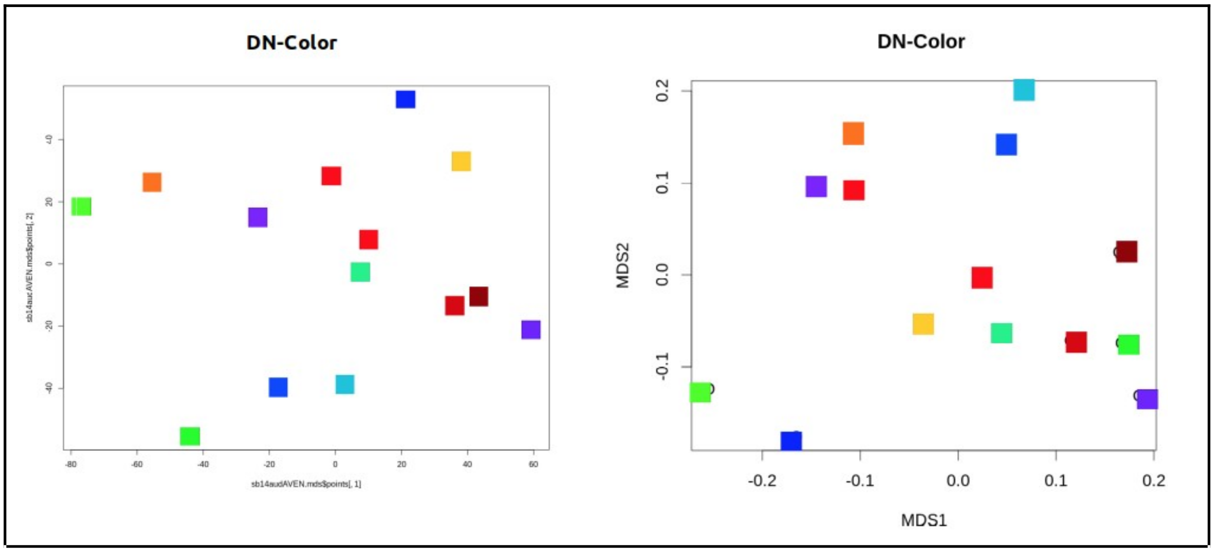


Figure S5. In panel a we show the nm-MDS of color stimuli decoded from Heshel's gyrus, producing a random set of coordinates for the presented wavelengths. In panel b, the neural distance matrix extracted from V4 but without boosting shows again a random set of coordinates for the presented wavelengths.

**(2) Letters**

To control for ROIs used for letters, we also used auditory cortex, and FEF, as well as PHG, areas that are less likely to produce the found SOI in extrastriate, although we might expect some other possible configurations including phonological similarity, or in PHG some similar SOI to ExtraStriate First in Figure S5a, we show the non-filtered BOLD with hierarchical clustering (Distance metric: euclidian, Grouping rule: WARD's). The dendrogram is color coded for the phrase structure in the

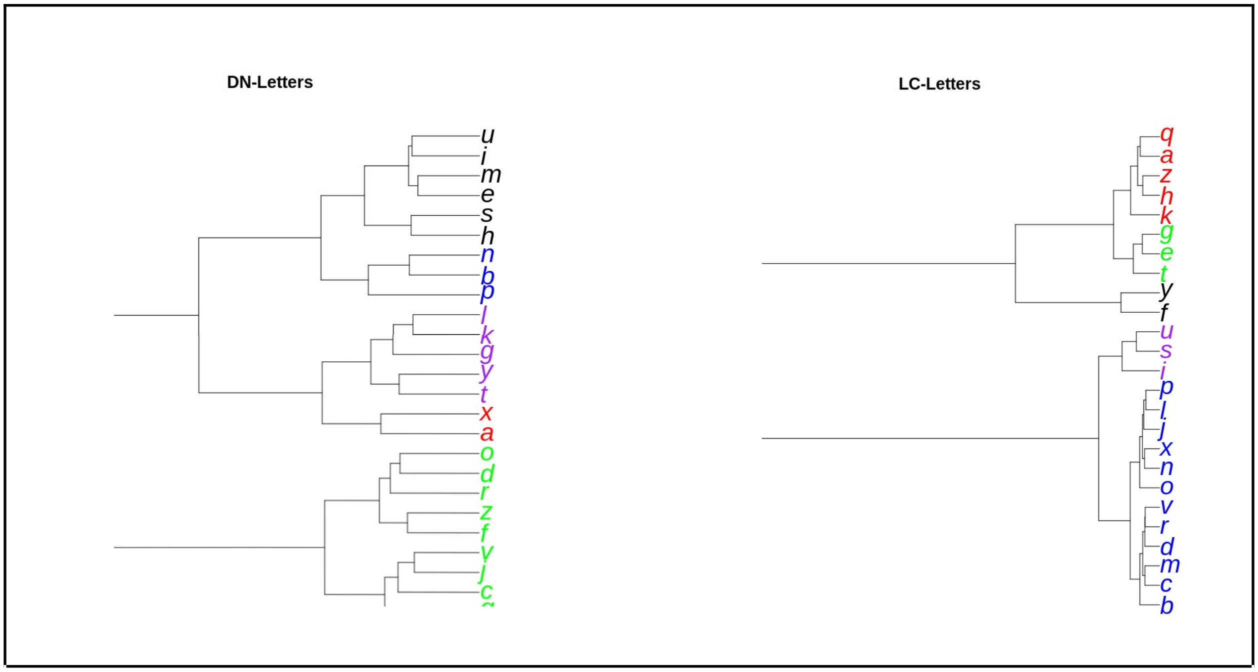


Figure S6. On the left we show the dendrogram recovered extracted from Heshel's gyrus (Auditory cortex) which does not recover the alphabet song phrasing, but does show some minimal phonological pairs (e.g., voicing bp, b-p, fricatives f-v). On the right we show the dendrogram without boosting for the same neural distance matrix in Figure 5 (main report) which again shows no clustering related to the Alphabet Song phrasing or sequential structure.

Alphabet song and it can be seen that no cluster that is derived from the raw BOLD data is consistent with the phrase structure of the alphabet mnemonic for learning letters. A second control test we ran was to decode from other non-visual cortical areas. In Figure S6b we show the result of pre-processed BOLD with the boosting filter, from auditory cortex (Heshel's gyrus), again the color code is based on the Alphabet Song phrase structure with no consistent clusters. It is interesting to note, that there are a few minimal phonological pairs (b:p; f:v) from auditory cortex decoding. We did similar tests with FEF, PHG also with no significant match to the SOI phrase structure.

**Random NDM tests**

To further control that the "figure-8" structure observed in the 3D nm-MDS configuration was not due to noise or nm-MDS artifact, we created a simulation by sampling from a Gaussian random variable . We first calculated the average mean and standard deviation of the subjects' decorrelated letter fMRI data, and then computed a distance matrix, and scaled it into 6D with nMDS. Resampling over six separate simulations, and plotting as one of the 3D projections of the 6D space, show (see Figure S7) that neither the sequential ordering of the letters, nor  the helical structure is observed indicating that we show the original decorrelated color data maintains its meaningful ordering and structure in the SOI found in 3D projections.

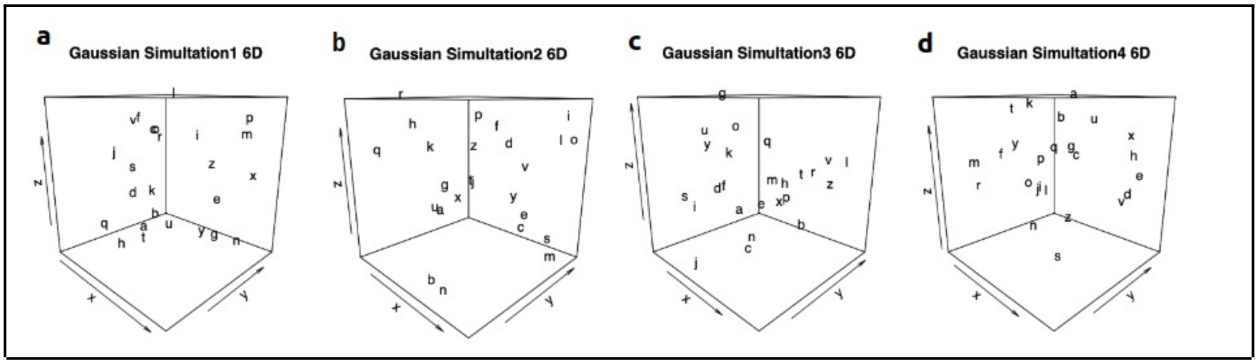


Figure S7. Shown here are 6D nMDS solutions for four gaussian random samples used to populate the “neural distance matrix” that was submitted to nMDS. None of the random cases shown (a, b, c, d)  exhibit the helical configuration, nor have any obvious alphabet sequential structure or other similar geometries across random samples.

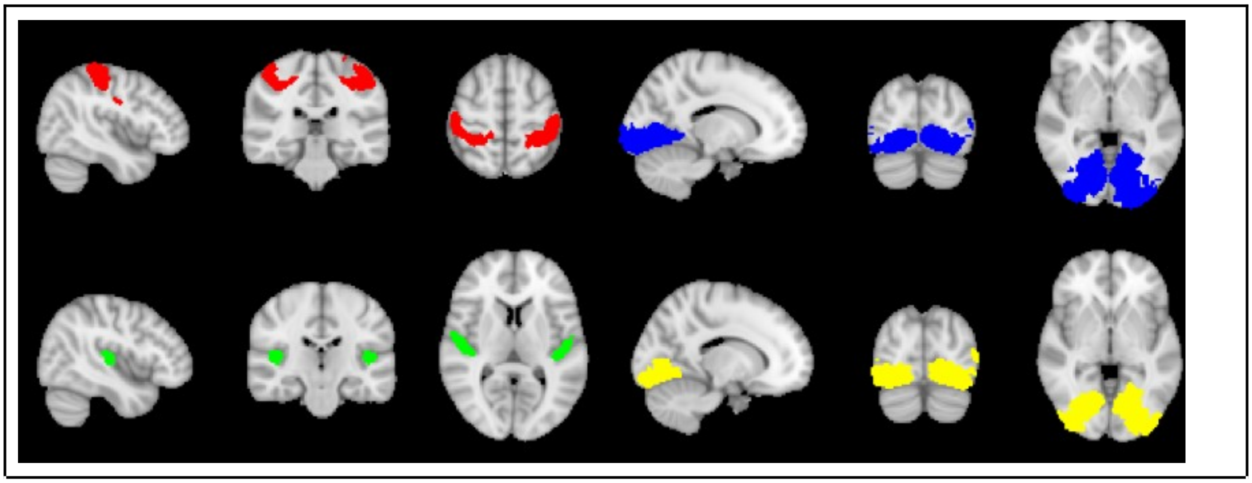

Figure S8.  Juelich atlas masks used for ROI time extraction. Somatosensory [red],  extrastriate (V1,V2,V3,V5)[blue], auditory cortex [green], and V4 [yellow]
